# Cell-free genome-wide transcriptomics through machine learning optimization

**DOI:** 10.1101/2025.06.04.657821

**Authors:** Léa Wagner, An Hoang, Olivier Rue, Olivier Delumeau, Valentin Loux, Jean-Loup Faulon, Matthieu Jules, Olivier Borkowski

## Abstract

Despite advances in transcriptomics, understanding of genome regulation remains limited by the complex interactions within living cells. To address this, we performed cell-free transcriptomics by developing a platform using an active learning workflow to explore over 1,000,000 buffer conditions. This enabled us to identify a buffer that increased mRNA yield by 20-fold, enabling cell-free transcriptomics. By employing increasingly complex conditions, our approach untangles the regulatory layers controlling genome expression.

## 2 Main

Lysate-based cell-free systems (CFS) are valuable platform for studying the living and elucidating molecular mechanisms^1–3^. Transcription and translation are restored in bacterial lysates by supplementing with a buffer containing ATP regeneration substrates, crowding agents, salts, and essential precursors such as nucleotides and amino acids. As such, CFS are simpler than living cells in terms of molecular interactions as they eliminate complex and interfering biological processes such as cell division and membrane-related functions. For example, previous experiments showed that production and metabolic burdens can be untangled using cell-free compared with *in vivo* measurements^4^. CFS provide an open platform ideally suited for protein production, high-throughput prototyping of genetic circuits and parts, biosensing biomanufacturing and gene expression analysis and modelling^5–7^.

Genome expression in living cells is routinely analyzed using RNA-seq^8^; however, low transcript yields in CFS have so far limited the application of this technique *in vitro*^9^. Here, we developed a general methodological pipeline that integrates active learning loops with high-throughput experimentation to optimize mRNA production in *Escherichia coli* BL21(DE3) CFS. This approach enabled transcript abundances sufficient for RNA-seq-based, genome-wide transcriptomic analysis. As proof of concept, we applied the pipeline to enhance RNA synthesis by T7 RNA polymerase (T7 RNAP) and performed *in vitro* transcriptomic profiling of the phage T7 genome.

To achieve mRNA abundances suitable for RNA-seq, T7 RNAP activity in CFS can be enhanced by optimizing the buffer composition. Machine learning has been successfully applied as an effective strategy for such optimization^10–13^. We selected 6 concentrations for each of the 8 components of our reference buffer (**Methods**), leading to more than one million possible compositions (**Fig. 1a**). To explore this large combinatorial space, we combined automated experimentation cycles with a machine learning algorithm based on Bayesian optimization with Gaussian processes (**Supplementary note 1, Supplementary fig. 1**), initiated by Latin Hypercube sampling (**Methods**), as illustrated in **Fig. 1b**. Automation of pipetting was performed using an acoustic liquid handler (Echo 550, Labcyte, USA). The algorithm was fed by mRNA abundance produced with each buffer composition. To quantify mRNA abundance in CFS, we used a reporter plasmid encoding deGFP and the malachite green aptamer, under control of a T7 promoter, enabling fluorescence-based readouts (**Fig. 1c**). This plasmid allows simultaneous monitoring of protein abundance and mRNA over time using a Synergy HTX plate reader (Synergy HTX, BioTek, USA). For each buffer, mRNA and protein yields were calculated as the maximum fluorescence normalized to the reference buffer (see **Methods**). Over cycles, data refined the machine learning model to suggest new sets of buffer compositions to test (**Supplementary note 2, Supplementary fig. 2**).

**Fig. 1.**
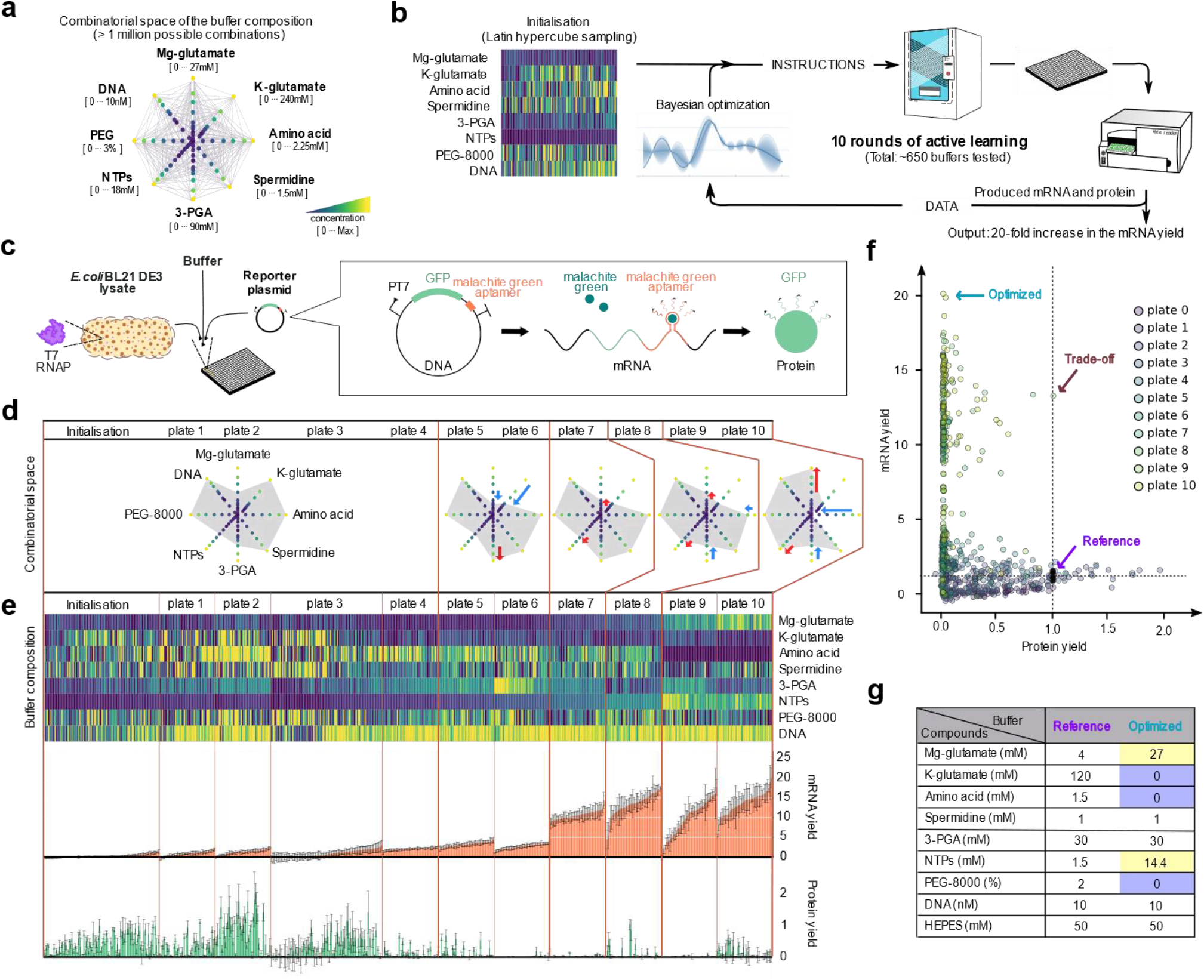
Active learning-based optimization of the cell-free buffer increases mRNA yield by 20-fold. **a**, Combinatorial space of buffer composition. The compounds and their maximum concentration range are indicated with the minimum concentration in blue and the maximum in yellow. **b**, Active learning pipeline for buffer optimization. The process was initiated with 100 buffer combinations, followed by 10 rounds of Bayesian optimization, resulting in a total of 653 unique buffer compositions tested. **c**, Schematic of the cell-free system setup: *E. coli* BL21 DE3 lysate is complemented with buffer and a reporter plasmid. Reporter plasmid encodes *degfp* and malachite green aptamer under the control of the T7 promoter for quantifying mRNA and protein abundances via fluorescence. All measurements were performed in 384 well-plates. **d**, Visualization of the combinatorial space explored on each plate, showing the distribution of buffer components across conditions. In each combinatorial space, red and blue arrows indicate increases and decreases, respectively, in the maximum concentration of a compound compared to the preceding space. **e**, Buffer optimization across 10 rounds of active learning. The heatmap gives the composition of the tested buffers, with the minimum concentration in blue and the maximum in yellow. Coral and green bar plots show respectively the median mRNA and protein yields for each buffer composition. Errors bars represent standard deviation between replicates (n=3 for plates 0-3; n=6 for plates 4-10). **f**, Scatter plot of protein yield versus mRNA yield for each buffer composition. Highlighted are the reference buffer (purple), trade-off buffer (brown), and mRNA-optimized buffer (blue). **g**, Composition of the reference and mRNA-optimized buffers. Compounds highlighted in yellow have increased concentrations compared to the reference, while those in blue have decreased concentrations. HEPES concentration was fixed during optimization.

**Fig. 1.**
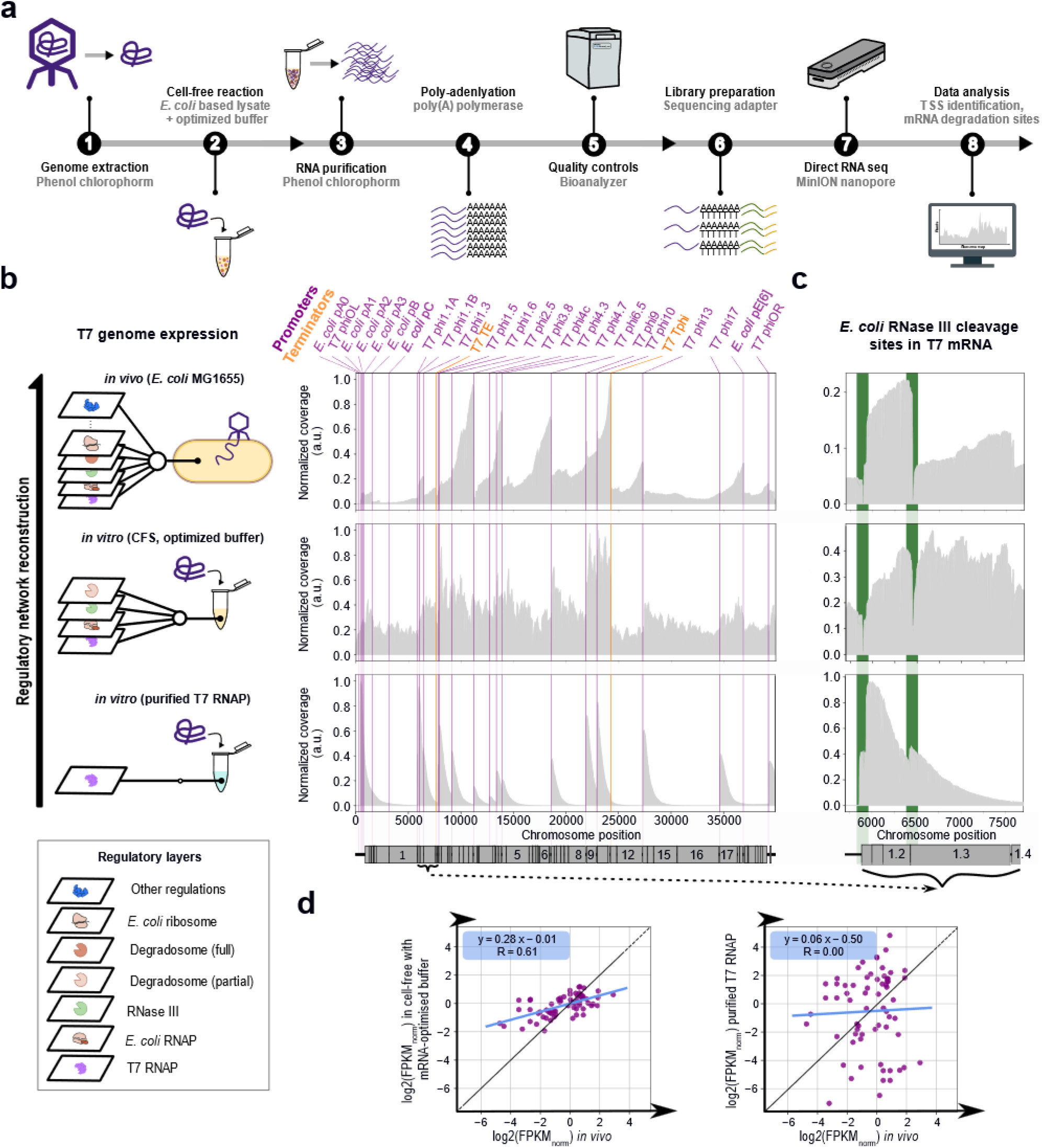
Comparative transcriptomics of T7 phage using cell-free system. **a**, Schematic of the cell-free transcriptomics workflow. **b**, Genome-wide expression profiles of T7 in various experimental conditions. On the left, schematic of the T7 genome expression *in vivo* and cell-free under optimized buffer condition, as well as with purified T7 RNA polymerase. On the right, sequencing depth (i.e., the number of times each base in the genome has been sequenced), normalized by the maximum value within each sample, is indicated by grey shaded areas. Promoters and terminators position are shown in purple and orange respectively. The position of the genes on the chromosome are represented in grey under the plots. **c**, Coverage plot highlighting genes 1.2 and 1.3, illustrating the activity of RNase III cleavage sites in T7 mRNA (in green). **d**, Correlation of log_2_-transformed median-normalized FPKM values between the *in vivo* and the other conditions. Gene expression values were normalized by the sample median FPKM to correct for global transcriptional differences and enable comparison of relative expression levels across conditions. In all panels, the blue line represents the linear regression and the black line indicates the identity line (y = x).

The concentration limits of each buffer component were constrained by the maximum cell-free reaction volume and the viscosity of stock solutions, which affected pipetting by the Echo machine. Over the iterations, the combinatorial space was progressively adjusted to maximize mRNA production without modifying the total reaction volume (10.5 µL), as shown by the grey area in **Fig 1d**. After 10 iterations, ∼650 buffer compositions were tested, leading to a 20-fold increase in mRNA yield with the optimized buffer (**Fig. 1e**). High Mg-glutamate, NTPs and DNA with low K-glutamate, and intermediate 3-PGA maximize mRNA abundance (**Fig. 1e, Supplementary note 3, Supplementary fig. 3**). Spermidine, amino acid and PEG-8000 seem to have no impact on mRNA abundance in cell-free (**Fig.1 e, Supplementary fig. 4**). Interestingly, the compositions which favored mRNA yields resulted in low protein yields (**Fig. 1f, Supplementary fig. 3**). Nevertheless, a trade-off between mRNA and protein yields was observed with one buffer composition, showing that higher mRNA abundance does not always correlate with lower protein abundance (**Fig. 1f, Supplementary note 4, Supplementary fig. 5**). The optimized buffer used for our cell-free RNA-seq analysis shows higher Mg-glutamate and NTPs, but lower K-glutamate, amino acids and PEG-8000 compared to the reference buffer (**Fig. 1g**).

We next expressed the T7 phage genome in the T7 optimized buffer for *in vitro* transcriptomic profiling. For comparison, the T7 genome was also expressed in two reference conditions: a minimalist *in vitro* transcription reaction using purified T7 RNAP in its commercial buffer, and a complex *in vivo* model involving infection of *E. coli* cells. Total RNA was purified, polyadenylated, and sequenced using direct RNA nanopore sequencing (MinION, Oxford Nanopore Technologies; **Fig. 1a**).

In the minimalist system, transcription was initiated from all T7-dependent promoters (**Fig. 1b, bottom row**). Coverage was skewed toward the 5′ end of transcripts, consistent with RNA accumulation in the absence of degradation. This makes the minimalist system an ideal proxy for comparing T7 RNAP-specific promoter activities in their native genomic context. Interestingly, promoters sharing the consensus sequence TAATACGACTCACTATAGGG showed position-dependent expression, underscoring the role of local genomic context (**Supplementary note 5, Supplementary fig. 6, Supplementary table 1**).

In CFS, transcripts from T7 RNAP-dependent promoters were detected as well (**Fig. 1b, middle row**). Uniform coverage along transcripts reflects a balance between transcription and degradation. Among degradation processes, T7 phage biology relies on mRNA maturation by RNase III^14^. Unlike the minimalist system, CFS restores T7 mRNA maturation, which is likely mediated by endogenous RNase III (**Fig. 1c, bottom and middle rows**).

*In vivo*, coverage was skewed toward the 3′ end of transcripts, indicating a different mRNA degradation than in CFS (**Fig. 1b, top row**). This may result from the absence of membrane-associated proteins, such as the primary endoribonuclease in *E. coli* RNase E^15,16^, in CFS^1^.

Overall, these three conditions represent increasing biological complexity: from purified T7 RNAP, to CFS, to *in vivo*, each step restoring additional layers of gene regulation. Our system can be expanded to non-model phages study^17^. Importantly, our work establishes a foundation for cell-free transcriptomics of complete bacterial genomes using endogenous RNA polymerases, providing a powerful platform for high-throughput *in vitro* analysis of bacterial gene regulation. This approach opens a way to access transcriptional landscape together with the regulation network of unculturable bacteria.

Additionally, the active learning workflow can be used to optimize both mRNA and protein yields, giving access to both transcriptomics and proteomics and allowing to investigate post-transcriptional mechanisms.

## 4 Methods

### 4.1 Strains and plasmids

Strain *E. coli* BL21 DE3 (B F^−^ *ompT gal dcm lon hsdS*_*B*_(*r*_*B*_^−^*m*_*B*_^−^) λ (DE3 [*lacI lacUV5*-*T7p07 ind1 sam7 nin5*]) [*malB*^+^]_K-12_(λ^S^)) was used to prepare the lysate ^18^. The plasmid PT7-deGFP-MGapt (4,176 bp) was a gift from Richard Murray (Addgene plasmid # 67741; http://n2t.net/addgene:67741 ; RRID: Addgene_67741) and transformed into chemically competent *E. coli* DH5α (F^−^ *endA1 glnV44 thi-1 recA1 relA1 gyrA96 deoR nupG purB20* φ80d*lacZ*ΔM15 Δ(*lacZYA-argF*)U169, hsdR17(*r*_*K*_^−^*m*_*K*_^+^), λ^−^).

### 4.2 Plasmid extraction and purification

The plasmid PT7-deGFP-MGapt was extracted from an overnight LB culture of *E. coli* DH5α using the QIAGEN Plasmid Mini Kit (Qiagen, ref 12123) following the manufacturer’s instructions. Since DNA preparation can significantly impact cell-free reactions^11,19^, we conducted all experiments using plasmid from the same initial stock. This stock was a combination of two batches prepared from 4 L and 3 L of the overnight LB culture, processed with 8 and 6 columns, respectively, and yielding approximately 4000 µg and 2400 µg of plasmid. The concentration of the stock was adjusted to 50nM (135.5 ng/µL) and aliquoted into 450 µL each.

### 4.3 Cell lysate preparation

The cell lysate preparation follows the protocol of Borkowski *et al*., which is based on the method developed by Sun *et al*.^11,20^. Due to malachite green’s sensitivity to DTT, all DTT was removed from the buffers^21^. Briefly, *E. coli* BL21 DE3 strain was cultured in rich 2xYT medium at 37°C with 1mM IPTG induction until the optical density reached 1.5-2.0. After two washes with DTT-free buffer S30A, each pellet was resuspended in a specified volume of buffer S30A based on its mass (1 g = 1 mL). The resuspended cells were then divided into 1 mL aliquots in 1.5 mL Eppendorf tubes. These tubes were sonicated using a Sonics VCX 750 Vibra Cell with 20% amplitude, following the sequence: 40 s ON – 60 s OFF – 40 s ON – 60 s OFF – 40 s ON. The remaining steps adhered to the protocol described by Sun *et al*., including a 1-hour run-off reaction at 37°C. The lysate was aliquoted into 2 mL Eppendorf tubes, with 1480 µL per tube, and stored at -70°C. The final lysate stock was prepared by combining 7 independently tested batches, totaling 32 L of culture and yielding 62 mL of lysate.

### 4.4 Cell-free reagents preparation

Cell-free reagents were prepared as previously described^20^. All products used and the final concentrations of our reagent stocks are listed in **Supplementary table 2**. Given that reagent preparation can significantly impact cell-free reactions^19^, we ensured that all our reagents came from aliquots of the same initial batch. The only exceptions were NTPs and 3-PGA, as additional material was needed during the study. We observed no effect on our controls when the NTPs and 3-PGA were renewed (**Supplementary fig. 7**).

To prepare the amino acid mixture, amino acids were grouped based on solubility and dissolved into three separate 36× stock solutions: water-soluble (neutral), acid-soluble, and base-soluble. Each amino acid was weighed according to its molecular weight to reach a final concentration of 54 mM per compound in 20 mL of stock solution. Water-soluble amino acids (alanine, arginine, glycine, histidine, lysine, proline, serine, threonine, and valine) were dissolved directly in ultrapure water. Acid-soluble amino acids (aspartic acid, asparagine, cystine, glutamine, glutamic acid, leucine, methionine, tryptophan, and tyrosine) were solubilized in 1 M HCl. Base-soluble amino acids (isoleucine and phenylalanine) were solubilized in 1 M KOH. Prior to use in cell-free experiments, the three stock solutions were mixed in equal volumes to yield a complete amino acid mixture.

### 4.5 Plasmid-based cell-free reaction

Reactions took place in 10.5 μL volumes at 30 °C in 384-well plate for 15h. To assemble the cell-free reactions, we used the Echo 550 liquid handler (Beckman, USA) to sequentially add a fixed amount of HEPES (50 mM), the variable buffer component, and finally, a fixed amount of malachite green (0.010 mM). The volume was then manually adjusted to 10.5 µL by adding a fixed amount of lysate (3.5 µL) and the appropriate volume of water using an automatic multi-channel pipette. The plate was immediately centrifuged at 1000 g for 1 minute at 4°C, sealed, and incubated in a plate reader preheated to 30°C.

### 4.6 Fluorescence measurements

We used a BioTek Synergy HTX plate reader (Agilent) to measure fluorescence in clear, flat-bottom 384-well plates (Greiner ref 781185). The plate was incubated for 15 hours at 30°C, with fluorescence measurements taken every 6 minutes and 7 seconds. The malachite green aptamer fluorescence was measured with excitation at 590±20 nm and emission at 635±20 nm, using a gain of 100 for plates 1-7 and a gain of 85 from plate 8 onward. GFP fluorescence was measured with excitation at 425±20 nm, emission at 510±20 nm, and a gain of 55. Fluorescence was detected from the top of the 384-well plate, which was sealed with clear sealing tape (Revivity ex. PerkinElmer #6050185 for plate 0 and ThermoFisher #232701 from plate 1 onward).

### 4.7 Definition of the reference buffer and combinatorial space

For *E. coli* lysate-based cell-free systems, a widely used method is the protocol established by Sun *et al*.^20^. In 2020, Borkowski *et al*. systematically optimized this buffer for protein production and identified five components that had minimal impact on protein yield^11^. In the present study, we adopted the Sun *et al*. buffer as a reference, omitting the non-essential components identified by Borkowski *et al*. resulting in the following composition: 4 mM Mg-glutamate, 120 mM K-glutamate, 1.5 mM amino acids, 1 mM spermidine, 30 mM 3-PGA, 1.5 mM NTPs, 2% PEG-8000, 50 mM HEPES. For the original combinatorial space, we defined six concentrations per component, anchoring the fifth at the reference level to enable exploration of four lower and one higher concentration.

### 4.8 Data analysis and normalization

The script “From_biotek_data_to_active_learning_vf.py” calculates a normalized value from the time-course malachite green aptamer and GFP fluorescence data obtained from the plate reader, enabling the comparison of mRNA and protein concentrations between buffers relative to a reference buffer. Briefly, the time-course fluorescence data were corrected well-by-well by subtracting the background fluorescence of the plastic, which was measured by reading the empty plate in the plate reader prior to filling it. For each well, the maximum fluorescence value was selected to calculate the yield as following:

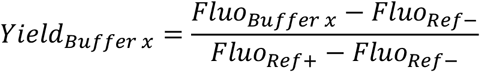

Where:

**Fluo**_**buffer_x**_ : For malachite green fluorescence, this is the maximum fluorescence value over time in each buffer composition. For GFP fluorescence data, it is the value after 15 hours, corresponding to the maximum fluorescence value in each buffer composition.

**Fluo**_**ref+**_ : The fluorescence (malachite or GFP) in the reference buffer containing 10 nM of DNA. **Fluo**_**ref-**_ : The fluorescence (red or green) with the reference buffer without DNA, representing the autofluorescence of the cell-free reaction, which is primarily due to the lysate autofluorescence.

Each yield represents the median of multiple replicates (3 for plates 0 to 3, and 6 for plates 4 to 10).

Starting from plate 7, the active learning process successfully identified optimal buffers for mRNA production, resulting in a significant increase in malachite green fluorescence. This increase required us to adjust a parameter—the gain—on the plate reader. Due to this new gain setting, the reference buffer with DNA described above could no longer be distinguished from the background noise. Therefore, we adjusted the yield calculation using an alternative reference buffer (P5_Buffer 21) that was previously tested in plates 5, 6, and 7:

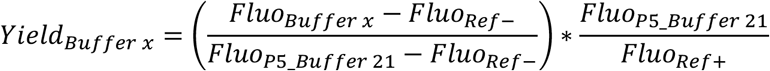

Where the ratio 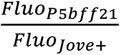 was fixed at 1.586, which is the median value of this ratio obtained from plates 5, 6, and 7. The composition of P5_Buffer 21 is: 2.01 mM Mg-glutamate, 40 mM K-glutamate, 2.25 mM amino acids, 0.50 mM spermidine, 45 mM 3-PGA, 3.74 mM NTPs, 2.49% PEG-8000, 50mM HEPES and 8.3 nM DNA.

To ensure that our normalization was sufficient for comparing data between plates, we measured a collection of control buffers on each plate and compared them across plates. The list of controls was iteratively updated over time to include buffers with increasingly higher fluorescence levels, reflecting the progression of active learning towards the optimal buffers. These controls validated that our normalization effectively corrected plate-to-plate variations, such as the change in plate sealer between plates 0 and 1 and the change in gain between plates 7 and 8 (**Supplementary fig. 8**).

### 4.9 Initialization of the active learning loop by Latin Hypercube sampling

To initiate the active learning loop, the initial 100 experiments were selected using Latin Hypercube Sampling to ensure good coverage of the search space. LHS assumes the independence of input variables (X). For each variable, the cumulative distribution function (based on a uniform distribution in our case) is divided into *n* equally sized partitions. A random value is then selected from each partition. These sampled values are combined across all variables to form *n* unique samples of X. One advantage of LHS arises when the output (Y, yield) is dominated by only a few components of X (cell-free mix components). This method ensures that these components are fully represented in a stratified manner, regardless of which components turn out to be important^22^. LHS is implemented by the SAMPLER module of the *icfree-ml* package (https://github.com/brsynth/icfree-ml).

### 4.10 Active learning modeling

The script “active_learning_bayesian_style_vf.py” generates a list of buffer compositions to test in the next plate according to active learning. Yield optimization was performed using a Bayesian optimization active learning approach, with a Gaussian Process (GP) model to probabilistically estimate the relationship between component concentrations and yield. The GP model was chosen for its robustness with small datasets and ability to capture predictions and uncertainties, outperforming more complex models prone to overfitting (**Supplementary note 1, Supplementary fig. 1**). Optimization was driven by the Upper Confidence Bound (UCB) acquisition function, with manual adjustments to the exploration-exploitation balance at each iteration to efficiently target high-potential regions of the search space. In each active learning cycle, the model provided a set of experiments in CSV format for execution using an Echo liquid handler.

### 4.11 Automated workflow

Modelling, experimental plate design, and translation into Echo instructions were automated using the LEARNER, DESIGNER, and INSTRUCTOR modules from the *icfree-ml* package (https://github.com/brsynth/icfree-ml), further described in Hérisson *et al*.^23^.

### 4.12 Amplification of T7 phage

A fresh colony of *E. coli* MG1655 grown on LB agar was used to inoculate a 5 mL overnight culture in Lennox rich medium, incubated at 37 °C with shaking at 200 rpm. The next day, the culture was diluted 1:100 into 5 mL of fresh Lennox medium supplemented with 10 mM MgSO_4_ and grown to an optical density at 600 nm (OD_600_) of 0.2. Two 1 mL aliquots were collected; one was infected with T7 phage at a multiplicity of infection (MOI) of 10^−4^ (i.e., 2×10^−4^ pfu per cell). After 10 min of adsorption at room temperature, both aliquots were diluted into separate 250 mL Erlenmeyer flasks containing 50 mL of prewarmed Lennox supplemented with 10 mM MgSO_4_. The uninfected culture served as a parallel control to monitor OD_600_, allowing an estimate of bacterial density in the infected culture during lysis. Once complete lysis was observed, the infected culture was centrifuged at 6,000 rpm for 10 min, and the supernatant was filtered through a 0.2 µm membrane. The resulting lysate was stored at 4°C and titrated.

### 4.13 Titration of T7 phage stock

A single colony of *E. coli* MG1655 was used to inoculate 3 mL of rich 2×YTP medium and grown overnight at 37°C with shaking at 200 rpm. The next day, 2 mL of fresh 2×YTP medium supplemented with 10 mM MgSO_4_ was inoculated with a 1:10 dilution of the overnight culture and incubated under the same conditions until exponential phase. In parallel, serial 1:100 dilutions of the T7 phage stock (to be quantified) were prepared in 1 mL of SM buffer (50 mM Tris-HCl pH 7.5, 100 mM NaCl, 10 mM MgSO_4_) to generate 10^− 1^, 10^− 3^, 10^−5^, 10^−7^, and 10^−9^ dilutions.

A soft LB-agar medium (7% agar, 10 mM MgSO_4_) was melted and kept at 50°C. For each phage dilution, 100 µL of the exponential-phase bacterial culture was spotted on one side of an empty Petri dish, and 100 µL of the corresponding phage dilution was spotted on the opposite side. The melted LB-agar was then poured into the plate, allowing the two drops to merge. Plates were gently rocked back and forth to ensure homogeneous distribution and left at room temperature until the agar solidified. After overnight incubation at 37°C, the phage titer was estimated by counting lysis plaques and accounting for the dilution factor, expressed in plaque-forming units per milliliter (pfu/mL).

### 4.14 Purification of T7 phage genome

Phage T7 were isolated using 50 ml of *E. coli* MG1655 growing in Lennox medium. Then, we centrifuged and filtered (0.2 µM) the lysed bacterial culture to remove debris. Phages are then concentrated by adding NaCl (0.5 M) and PEG-8000 (10%) and incubating overnight at 4°C. After centrifugation, the pellet is resuspended in SM buffer, and RNase A and DNaseI are added to remove RNA and digest free DNA. Following a 30-minute incubation, DNaseI is inactivated with EDTA. Phage DNA is then extracted using phenol-chloroform and precipitated with potassium acetate and ethanol. The DNA is washed, resuspended in water, and quantified using a Qubit fluorometer.

### 4.15 Genome-based expression and RNA purification in the minimalist system

The T7 genome was expressed using either purified T7 RNA polymerase (TranscriptAid T7 High Yield Transcription Kit, Thermo Fisher #K0441) in a 20 µL reaction or purified *E. coli* RNA polymerase (Holoenzyme, NEB M0551). After a 2-hour incubation at 37°C, the RNA was treated with DNase I (2 µL) for 15 minutes at 37°C, followed by 2 µL 0.5 M EDTA (pH 8.0) and incubation at 65°C for 10 minutes to stop the reaction. RNA was then purified using the Monarch RNA Cleanup Kit (NEB T2050), polyadenylated (see section 4.18), and sequenced (see section 4.19).

### 4.16 Genome-based expression and RNA purification from cell-free reactions

The genome-based cell-free reactions were performed in the same way as the plasmid-based cell-free reactions, except that the DNA was not sent by the ECHO but added manually just before the lysate. To prevent digestion of the linear T7 genome in cell-free, a Gibson reaction was performed taking advantage of the 160bp repeated sequence naturally present in T7 genome (NEBuilder HiFi DNA Assembly Master Mix, NEB #E2621). DNA was dialyzed for 30min using 0.025 µM membrane (Millipore #VSWP01300) before use. RNA extraction from cell-free reactions was performed after 3h of reaction, which corresponds to the time to reach peak RNA concentration in the optimized buffer (P10_Buffer 9).

RNA was purified from cell-free reaction by phenol/chloroform extraction as described previously^24^. Briefly, the samples were extracted twice with an equal volume of acid phenol/chloroform/isoamyl alcohol (25:24:1, [pH 4.5]) and once with chloroform/isoamyl alcohol (24:1). After adding 1/10 volume of 3 M sodium acetate (pH 5.2), RNA was precipitated with isopropanol, washed with 70% ethanol and dissolved in 100 μl of RNase free water. The samples were then DNase-treated using the RNase-Free DNase Set (Qiagen) and purified using the RNA Clean-Up and Concentration Micro Kit (Norgen). The RNA concentration was measured using a NanoDrop spectrophotometer; the quality of the RNA preparations was assessed by means of the Agilent 2100 Bioanalyzer according to the manufacturer’s instructions.

### 4.17 Infection and RNA purification for *in vivo* experiment

An overnight culture of *E. coli* MG1655 was prepared in 2 mL of rich 2×YTP medium supplemented with 10 mM MgSO_4_. The culture was diluted 1:400 into 150 mL of fresh 2×YTP medium and incubated at 37 °C with shaking at 200 rpm until the optical density at 600 nm (OD_600_) reached 0.5. A second 1:400 dilution was then used to inoculate 150 mL of 2×YTP for the main culture. Upon reaching OD_600_ = 0.3 (corresponding to approximately 1.5 × 10^8^ cells/mL), the culture was split in two, and one half was infected with T7 phage at a multiplicity of infection (MOI) of 1. Ten minutes post-infection—within the estimated duration of the T7 lytic cycle (5–12 min)— cells were harvested by centrifugation (3 min, 8,000 × g, 4 °C), and the pellet was flash-frozen in liquid nitrogen and stored at -70°C.

The next day, total RNA was isolated as described previously^24^. Briefly, bacterial cultures were mixed with one-half volume of frozen killing buffer (20 mM Tris-HCl, pH 7.5; 5 mM MgCl_2_; 20 mM NaN_3_), and cells were harvested by centrifugation at 4°C for 3 min. After discarding the supernatant, cell pellets were flash-frozen in liquid nitrogen and stored at –80 °C. For mechanical lysis, pellets were resuspended in 200 µL of ice-cold killing buffer and immediately transferred to a precooled Teflon disruption vessel containing liquid nitrogen. Samples were disrupted using a Mikro-Dismembrator S (Sartorius) for 2 min at 2,600 rpm. The resulting frozen powder was resuspended by pipetting in 4 mL of lysis buffer prewarmed to 50°C (4 M guanidine thiocyanate, 25 mM sodium acetate pH 5.2, 0.5% [wt/vol] N-laurylsarcosinate). Aliquots of 1 mL were transferred to microcentrifuge tubes and immediately flash-frozen in liquid nitrogen. Total RNA was purified by phenol–chloroform extraction, as described in the preceding section.

### 4.18 Polyadenylation of RNA

For direct RNA sequencing with Oxford Nanopore Technology (ONT), polyadenylation was performed following the ONT protocol (v4, March 2023) using *E. coli* Polya, Polymerase (NEB: M0276L). This version of the ONT protocol is provided as “3-poly-rna-ecoli-pap.pdf” and is available for download at: https://nanoporetech.com/document/extraction-method/3-poly-rna-ecoli-pap?format=versions. Briefly, RNA was incubated at 37°C for 1 minute with 5 U of Polya, Polymerase and 1 mM ATP, then the reaction was halted with EDTA. Polyadenylated RNA was purified using beads.

### 4.19 Long reads direct RNA sequencing with Oxford Nanopore Technology

Library preparation was carried out using the Direct RNA Sequencing Kit (SQK-RNA004, ONT) for sequencing on the Flow Cell RNA (FLO-MIN004RA, ONT) on a MinION Mk1B (ONT) device using the MinKNOW version 24.06.5 software. The ONT protocol used in this study was an early access version, which is provided as “direct-rna-sequencing-sqk-rna004-DRS_9195_v4_revB_20Sep2023-minion.pdf” and is available for download at: https://community.nanoporetech.com/attachments/11348/download. RNA Control Strand (RNACS), corresponding to the *S. cerevisiae* ENO2 gene (YHR174W), was added to all samples at a defined concentration as an external spike-in, following protocol recommendations.

### 4.20 RNA-seq data analysis

RNA-seq data analysis was performed using the Migale bioinformatics facility. The entire workflow of the analysis is presented in **Supplementary fig. 9**, detailing the names and versions of all bioinformatics tools used, the custom Python scripts developed for this study, and the intermediate files made available in the accompanying data repository (https://doi.org/10.57745/BHTM60). Code snippets and usage details are provided in the repository report.

#### Data preparation

Only reads from the “pass” folder generated by MinKNOW were used, corresponding to those passing the initial quality threshold (Q score above 8). Raw direct RNA sequencing reads were concatenated into a single fastq file per flow cell. To ensure compatibility with standard DNA-based bioinformatics tools, uracil (U) residues were converted to thymine (T).

#### Quality control and reads cleaning

Read quality was assessed using seqkit^25^, FastQC (Simon A., https://www.bioinformatics.babraham.ac.uk/projects/fastqc/(2010)), MutliQC^26^, NanoStat^27^ and NanoPlot^27^, enabling evaluation of read length distribution, GC content, quality score distributions, and overall sequencing performance. Reads were filtered using Chopper^27^, a tool optimized for long-read data, retaining only those with GC content between 30–65% and lengths below 40,000 bases, thus removing aberrant reads and sequencing artifacts. The total number of reads obtained for each flow cell is reported in **Supplementary fig. 10**.

#### Read mapping

As expected, reads mapped to three reference sequences: (i) the T7 genome, (ii) the *E. coli* BL21(DE3) genome, and (iii) the *S. cerevisiae* ENO2 gene (YHR174W), used as an external spike-in control. Although the lysate is expected to be free of *E. coli* genomic DNA and RNA, degradation products of *E. coli* ribosomes (16S and 23S rRNA) are expected to generate reads that map to the *E. coli* genome. The three references were concatenated into a single reference genome (3refs.fasta). To prevent redundancy, the T7 RNA polymerase gene was identified in the *E. coli* genome using BLAST^28^ and masked using BEDTools^29^. Reads were aligned to this composite reference using minimap2, with secondary and supplementary alignments disabled^30^.

#### Reference-specific read quantification

Using alignment flags in the resulting BAM files, the proportion of reads mapping to each of the three references was computed using samtools^31^ (**Supplementary fig. 10**).

#### Coverage analysis

From the BAM files, base-level coverage was extracted using samtools to generate a three-column text file (chromosome, position, depth), named “ID_vs_REF_coverage.txt”^31^.

#### FPKM quantification

Gene expression levels were estimated from the BAM alignments using BEDTools^29^. Output included transcript-level FPKM values stored in “gene_counts_ID_vs_REF.fpkm.txt”. For long-read sequencing, FPKM are calculated as:

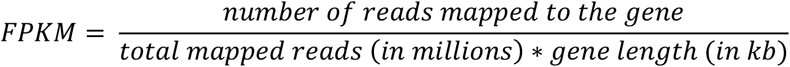

#### Data integration and visualization

Python scripts were used to integrate and visualize expression data. The script “create_tables.py” merged per-base coverage with FPKM values, producing two output files: “ID_vs_REF_coverage_by_position.tsv” and “ID_vs_REF_FPKM_by_gene.tsv”, summarizing base-level and gene-level data, respectively. These were used to generate coverage plots with overlaid gene expression levels, using the script “plot_coverage_FPKM.py” (**Supplementary note 6, Supplementary fig. 11**). The script “plot_correlation_FPKM.py” was used to compare FPKM values between flow cells.

## Acknowledgments

We thank Joan Herisson and Mahnaz Sabeti Azad for providing the ECHO code and helpful explanations; Patricia Lepage for access to computing resources; Paul Soudier for the amino acid mixture formulation; Jérôme Bonnet and Vincent Libis for their valuable advice throughout the project; the @BRIDGe platform (INRAE, Ile-de-France, Jouy-en-Josas) for providing access to the Bioanalyzer; Marie-Agnès Petit and Inès Doublier for support with phage experiments; Sophie Pollet and Mathilde Filipe-Ferreira for helpful advice on MinION sequencing; and Elena Bidnenko for guidance on RNA purification protocols. We are grateful to the INRAE MIGALE bioinformatics facility (MIGALE, INRAE, 2020. Migale bioinformatics Facility, https://doi.org/10.15454/1.5572390655343293E12) for providing help and computing and storage resources.

## 6 Author contributions

L.W., M.J. and O.B. designed the study. L.W. and O.B. conducted cell-free experiments. L.W. and A.H. developed the active learning loop. JL. F supervised the cell-free automation. L.W. developed the RNA-seq workflow and performed experiments. L.W., O.R. and V.L. analyzed the RNA-seq data. O.D. supervised the setting up of the experiments using malachite green and in particular shared his knowledge of the impact of DTT. O.B. and L.W. designed figures with inputs from M.J. and O.D. L.W., M.J. and O.B. wrote the manuscript. All authors reviewed and approved the final version of the manuscript.

## 7 Competing interests

All authors declare no competing interests.

## 8 Funding

This work has been supported by the ANR program, RNADarkMatter “ANR-24-CE44-4467”, ANR-21-ESRE-0021, by France 2030 program “ANR-11-IDEX-0003”. L.W. acknowledges a 3-year Ph.D. grant from INRAE, France. JL. F and A.N. H acknowledges the ANR-22-PEBB-0008 (PEPR B-BEST France 2030 program) and UE HORIZON BIOS program (grant number 101070281).

## Notes

### Competing Interest Statement

The authors have declared no competing interest.

## References

1. Wagner L, Jules M, Borkowski O. What remains from living cells in bacterial lysate-based cell-free systems. Computational and Structural Biotechnology Journal. 2023 Jan 1;21:3173–82.

2. Giaveri S, Bohra N, Diehl C, et al. Integrated translation and metabolism in a partially self-synthesizing biochemical network. Science. 2024 Jul 12;385(6705):174–8.

3. Kosaka Y, Miyawaki Y, Mori M, et al. Autonomous ribosome biogenesis in vitro. Nat Commun. 2025 Jan 8;16(1):514.

4. Borkowski O, Bricio C, Murgiano M, et al. Cell-free prediction of protein expression costs for growing cells. Nat Commun. 2018 Apr 13;9(1):1457.

5. Siegal-Gaskins D, Tuza ZA, Kim J, et al. Gene circuit performance characterization and resource usage in a cell-free “breadboard.” ACS Synth Biol. 2014 Jun 20;3(6):416–25.

6. Silverman AD, Karim AS, Jewett MC. Cell-free gene expression: an expanded repertoire of applications. Nat Rev Genet. 2020 Mar;21(3):151–70.

7. Hunt AC, Rasor BJ, Seki K, et al. Cell-free gene expression: methods and applications. Chem Rev. 2025 Jan 8;125(1):91–149.

8. Zhu C, Preissl S, Ren B. Single-cell multimodal omics: the power of many. Nat Methods. 2020 Jan;17(1):11–4.

9. Fujiwara K, Sawamura T, Niwa T, et al. In vitro transcription–translation using bacterial genome as a template to reconstitute intracellular profile. Nucleic Acids Res. 2017 Nov 2;45(19):11449–58.

10. Caschera F, Bedau MA, Buchanan A, et al. Coping with complexity: machine learning optimization of cell-free protein synthesis. Biotechnology and Bioengineering. 2011;108(9):2218–28.

11. Borkowski O, Koch M, Zettor A, et al. Large scale active-learning-guided exploration for in vitro protein production optimization. Nat Commun. 2020 Apr 20;11(1):1872.

12. Morini L, Sakai A, Vibhute MA, et al. Leveraging active learning to establish efficient in vitro transcription and translation from bacterial chromosomal DNA. ACS Omega. 2024 Apr 30;9(17):19227–35.

13. Wang H, Fu T, Du Y, et al. Scientific discovery in the age of artificial intelligence. Nature. 2023 Aug;620(7972):47–60.

14. Dunn JJ, Studier FW. T7 early RNAs and Escherichia coli ribosomal RNAs are cut from large precursor RNAs in vivo by ribonuclease III. Proceedings of the National Academy of Sciences. 1973 Dec;70(12):3296–300.

15. Mudd EA, Krisch HM, Higgins CF. RNase E, an endoribonuclease, has a general role in the chemical decay of Escherichia coli mRNA: evidence that rne and ams are the same genetic locus. Mol Microbiol. 1990 Dec;4(12):2127–35.

16. Khemici V, Poljak L, Luisi BF, et al. The RNase E of Escherichia coli is a membrane-binding protein. Mol Microbiol. 2008 Nov 1;70(4):799–813.

17. Putzeys L, Wicke L, Brandão A, et al. Exploring the transcriptional landscape of phage–host interactions using novel high-throughput approaches. Current Opinion in Microbiology. 2024 Feb 1;77:102419.

## References

18. Studier FW, Moffatt BA. Use of bacteriophage T7 RNA polymerase to direct selective high-level expression of cloned genes. J Mol Biol. 1986 May 5;189(1):113–30.

19. Cole SD, Beabout K, Turner KB, et al. Quantification of interlaboratory cell-free protein synthesis variability. ACS Synth Biol. 2019 Sep 20;8(9):2080–91.

20. Sun ZZ, Hayes CA, Shin J, et al. Protocols for implementing an Escherichia coli based TX-TL cell-free expression system for synthetic biology. J Vis Exp. 2013 Sep 16;(79):e50762.

21. Jurado Z, Murray RM. Impact of chemical dynamics of commercial PURE systems on malachite green aptamer fluorescence. ACS Synth Biol. 2024 Oct 18;13(10):3109–18.

22. McKay MD, Beckman RJ, Conover WJ. A comparison of three methods for selecting values of input variables in the analysis of output from a computer code. Technometrics. 1979;21(2):239–45.

23. Hérisson J, Hoang AN, El-Sawah A, et al. Operate a cell-free biofoundry using large language models. Preprint at: https://www.biorxiv.org/content/10.1101/2024.10.28.619828v1 (2025)

24. Nicolas P, Mäder U, Dervyn E, et al. Condition-dependent transcriptome reveals high-level regulatory architecture in Bacillus subtilis. Science. 2012 Mar 2;335(6072):1103–6.

25. Shen W, Le S, Li Y, Hu F. SeqKit: a cross-platform and ultrafast toolkit for FASTA/Q file manipulation. PLOS ONE. 2016 Oct 5;11(10):e0163962.

26. Ewels P, Magnusson M, Lundin S, et al. MultiQC: summarize analysis results for multiple tools and samples in a single report. Bioinformatics. 2016 Oct 1;32(19):3047–8.

27. De Coster W, Rademakers R. NanoPack2: population-scale evaluation of long-read sequencing data. Bioinformatics. 2023 May 1;39(5):btad311.

28. Altschul SF, Gish W, Miller W, et al. Basic local alignment search tool. Journal of Molecular Biology. 1990 Oct 5;215(3):403–10.

29. Quinlan AR, Hall IM. BEDTools: a flexible suite of utilities for comparing genomic features. Bioinformatics. 2010 Mar 15;26(6):841–2.

30. Li H. Minimap2: pairwise alignment for nucleotide sequences. Bioinformatics. 2018 Sep 15;34(18):3094–100.

31. Li H, Handsaker B, Wysoker A, Fennell T, Ruan J, Homer N, et al. The sequence alignment/map format and SAMtools. Bioinformatics. 2009 Aug 15;25(16):2078–9.

